# Lysosomes are Required for Early Dorsal Signaling in the *Xenopus* Embryo

**DOI:** 10.1101/2022.01.19.476939

**Authors:** Nydia Tejeda-Muñoz, Edward M. De Robertis

**Author notes:** **Author contributions:** N.T-M. and E.M.D.R. designed research, performed research, analyzed data, and wrote the paper.

## Abstract

Lysosomes are the digestive center of the cell and play important roles in human disease, including cancer. Previous work has suggested that late endosomes, also known as multivesicular bodies (MVBs), and lysosomes are essential for canonical Wnt pathway signaling. Sequestration of Glycogen Synthase 3 (GSK3) and of β□catenin destruction complex components in MVBs is required for sustained canonical Wnt signaling. Little is known about the role of lysosomes during early development. In the *Xenopus* egg, a Wnt-like cytoplasmic determinant signal initiates formation of the body axis following a cortical rotation triggered by sperm entry. Here we report that cathepsin D was activated in lysosomes specifically on the dorsal marginal zone of the embryo at 64-cell stage, long before zygotic transcription starts. Expansion of the multivesicular body (MVB) compartment with low-dose Hydroxychloroquine (HCQ) greatly potentiated the dorsalizing effects of the Wnt agonist Lithium chloride (LiCl) in embryos, and this effect required macropinocytosis. Formation of the dorsal axis required lysosomes, as indicated by brief treatments with the vacuolar ATPase (V-ATPase) Bafilomycin A1 inhibitor at the 32-cell stage. Inhibiting the MVB-forming machinery with a dominant-negative point mutation in Vacuolar Protein Sorting 4 (Vps4-EQ) also interfered with the endogenous dorsal axis. The Wnt-like activity of the dorsal cytoplasmic determinant Huluwa (Hwa), and that of microinjected xWnt8 mRNA, also required lysosome acidification and the MVB-forming machinery. We conclude that lysosome function is essential for early dorsal axis development in *Xenopus*. The results highlight the intertwining between membrane trafficking, lysosomes, and vertebrate axis formation.

**Significance:** The dorsal axis of the vertebrate Xenopus embryo is established by an early Wnt signal generated by a rotation of the cortex of the egg towards the opposite side of the sperm entry point. In this study, we report that lysosomal Cathepsin D becomes activated on the dorsal marginal zone of the embryo already at the 64-cell stage, and that this asymmetry is enhanced by increasing Wnt signaling levels. We present experiments showing that lysosome activity, macropinocytosis, and multivesicular body formation are required for the dorsal signal provided maternally in the egg, and for twinning by microinjected *huluwa* and *Wnt8* mRNA. The results indicate that the cell biology of lysosomes plays a fundamental role in vertebrate development.

## INTRODUCTION

In *Xenopus* development, formation of the Spemann organizer and subsequent axial development is controlled by a dorsal Wnt-like signal resulting from a microtubule-driven rotation of egg cortical cytoplasm towards the opposite side of the sperm entry point (1, 2). This signal is known to involve the displacement of cytoplasmic membrane vesicles located on the vegetal pole that contain Dishevelled (Dvl) (3, 4). Recently, it has been proposed that the zebrafish maternal determinant is a novel transmembrane protein called Huluwa (Hwa), which causes degradation of the Wnt pathway component Axin1 in dorsal blastomeres (5). While all transcriptional effects of Wnt signaling are mediated by the stabilization of β-catenin (6, 7), recent work has revealed that membrane trafficking is also essential for Wnt signaling (8, 9, 10). The inhibition of glycogen synthase 3 kinase (GSK3) after binding to the Wnt receptors Lrp6 and Frizzled triggers macropinocytosis, an actin-driven cell drinking process (11, 12) by which extracellular proteins are engulfed and trafficked to lysosomes for degradation (12, 13). The increased membrane trafficking leads to the formation of multivesicular bodies (MVBs) that sequester GSK3 and Axin1, resulting a marked increase in lysosomal acidification and activation of lysosomal enzymes (13).

Here we report that the active form of the lysosomal enzyme cathepsin D localizes to dorsal cells in 64-cell *Xenopus* embryos, and that this activation is greatly increased by microinjection of *xWnt8* mRNA or the GSK3 inhibitor Lithium chloride (LiCl). Expanding the MBV late endosomal compartment by microinjecting low-dose Hydroxychloroquine (HCQ) potentiated dorsal signaling by LiCl. This effect was blocked by microinjection of 5-N-ethyl-N-isopropylamiloride (EIPA), a macropinocytosis inhibitor. In the intact embryo, Spemann organizer formation and central nervous system (CNS) induction were inhibited by short treatment with the vacuolar ATPase (V-ATPase) inhibitor Bafilomycin A1 (Baf) or by interfering with MVB formation. Signaling by microinjected *xHwa* mRNA was blocked by Baf treatment at the 32-cell stage, co-injection with EIPA, or inhibitors of the ESCRT (Endosomal Sorting Complexes Required for Transport) machinery. We conclude that lysosomes play an essential role in the initiation of axis formation by the dorsal cytoplasmic determinant.

## RESULTS

### Activated Lysosomes in the Dorsal Side at 64-cell Stage

Cortical rotation in *Xenopus* results in a less pigmented dorsal crescent that marks the future dorsal side during early development (Fig. 1*A, A’*) (14). Embryos at 64-cell stage were fixed, bisected, and stained with Sir-Lysosome, a very useful reagent that specifically labels active lysosomes by binding to the activated, cleaved form of lysosomal cathepsin D via a pepstatin A peptide (13, 15). An enrichment in active lysosomes was observed in the dorsal marginal cells of uninjected control embryos (Fig. 1*B, B’*). This asymmetry was strongly enhanced by microinjection of *xWnt8* mRNA (16) or LiCl (17) (Fig. 1*C*-*D’*). In addition, LiCl microinjection caused a striking stabilization of the V-ATPase subunit V0a3 (18), particularly on the dorsal side of the embryo (*SI Appendix*, Fig. S1*A*-*B’* available online). The stabilization of V-ATPase, the enzyme that acidifies the endolysosomal pathway and controls membrane trafficking, is consistent with previous observations that GSK3 inhibition increases macropinocytosis and subsequent lysosomal acidification (13). The results indicate that lysosomes are activated in the dorsal side of the embryo at early cleavage stages, long before zygotic transcription is activated.

**Fig. 1.**
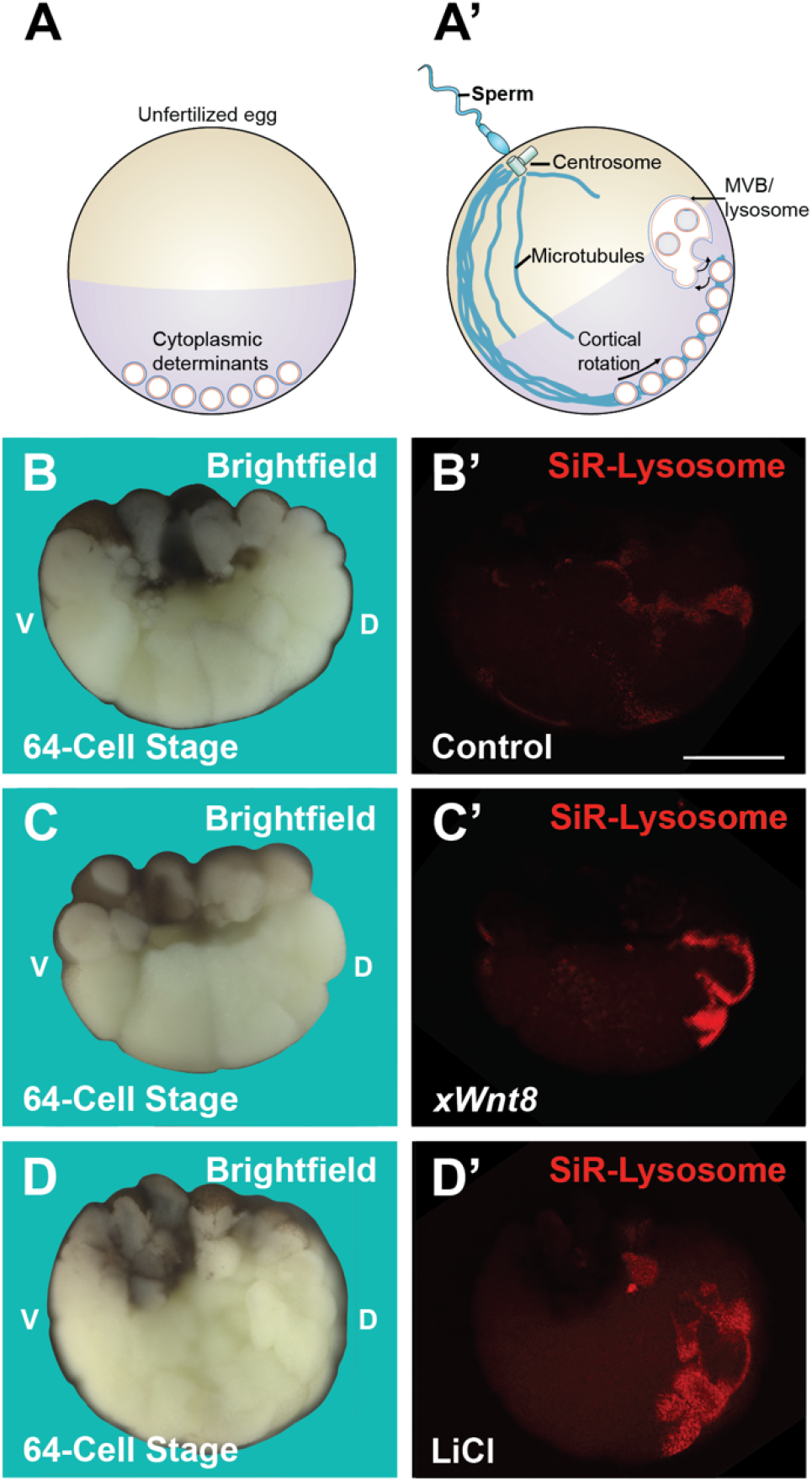
Lysosomes are Activated on the Dorsal Side of 64-cell Stage Embryos and this Asymmetry is Increased by *xWnt8* mRNA or LiCl Microinjection. (*A*) The unfertilized egg contains cytoplasmic determinants indicated by membrane-bounded organelles that contain Dvl. (*A’*) Sperm entry introduces the centrosome which nucleates microtubules that drive cortical rotation of cytoplasmic determinats. The hypothesis tested here is that MVBs and lysosomes are activated by the maternal Wnt-like signal. (*B* and *B’*) SiR-lysosome fluorophore stains activated cathepsin D in dorsal cells at 64-cell stage. (*C’*-*D’*) Microinjection of 2 pg *xWnt8* mRNA or 4 nl of 300 mM LiCl into the vegetal pole at 4 cells greatly increases active lysosomes. Numbers of embryos analysed were as follows B=31, 100%; C=29, 93.2%; D=32, about 90% with dorsal signal, 5 independent experiments. Scale bars, 500 μm. See also *SI Appendix*, Fig. S1.

### Low-dose Hydroxychloroquine Increases Wnt and LiCl Signaling

We have shown that low doses of the lysosomotropic antimalarial drug Chloroquine (CQ) can increase Wnt signaling 2-3-fold by expanding the MVB compartment and facilitating the sequestration of GSK3 and Axin1 in MVBs (19). We now report that its derivative Hydroxychloroquine (HCQ) is more effective than CQ, increasing Wnt3a-induced β-catenin activated reporter (BAR) expression (20) up to 34 times in cultured mammalian cells (Fig. 2A). In the absence of Wnt, HCQ had no stimulatory effect (Fig. 2A). At high concentrations HCQ, like CQ, had the opposite effect, inhibiting Wnt by alkalinizing the endosomal compartment (*SI Appendix*, Fig. S2*A*-*G*). In the presence of Wnt, low-dose HCQ enhanced Wnt signaling, causing stabilization of CD63 (Fig. S2*H*-*L*), a marker of MVB intraluminal vesicles (21). The transcriptional β-catenin signal triggered by the GSK3 inhibitor LiCl was also enhanced by low-dose HCQ, although the optimal concentration level was different from that of Wnt (Fig. 2B).

**Fig. 2.**
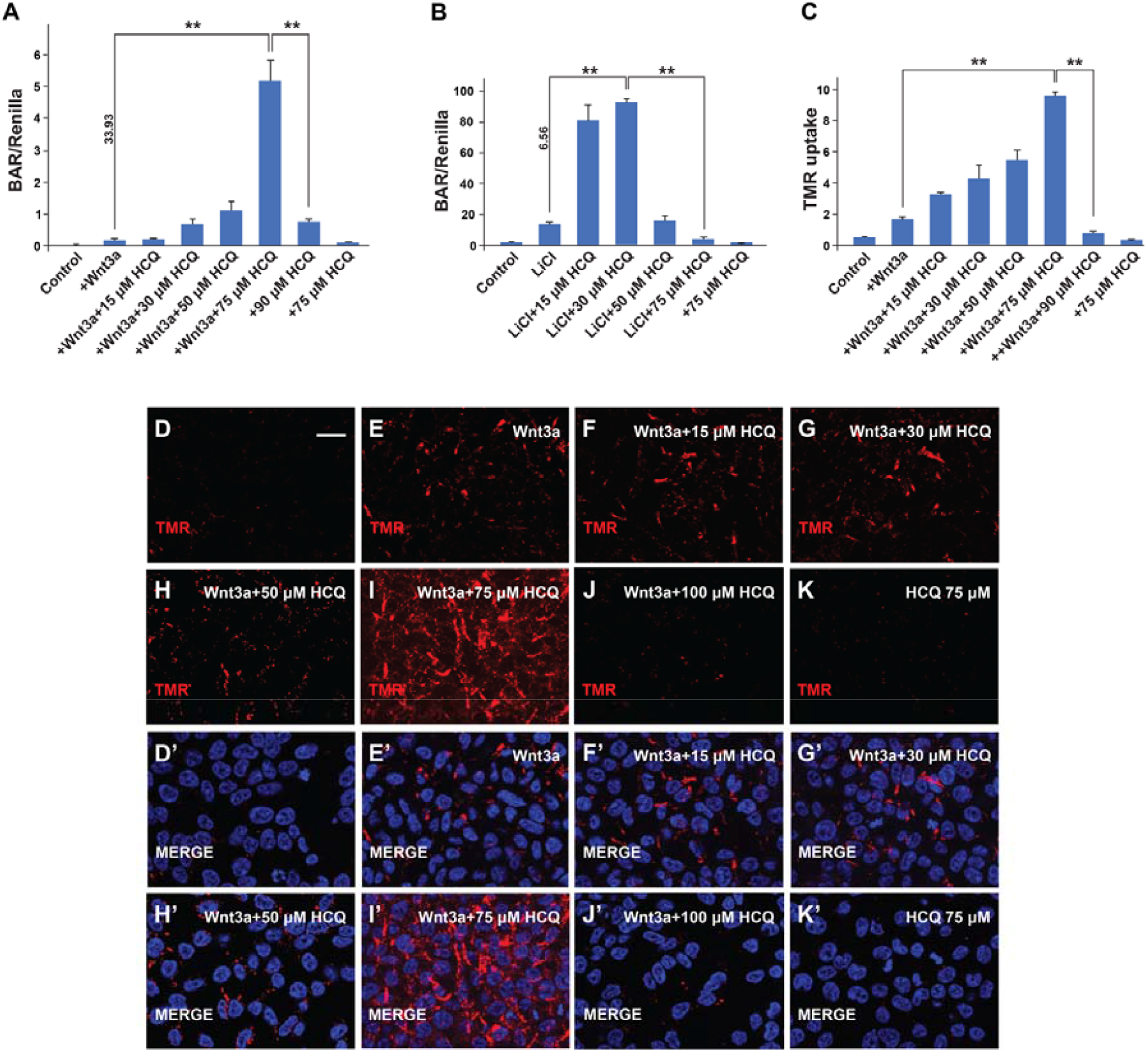
Wnt3a, LiCl Transcriptional Activity and Macropinocytosis, are Potentiated by Low-dose Hydroxychloroquine. All assays were performed in HEK293-BR cells that contain stably integrated BAR and Renilla reporter genes. (*A*) Wnt3a treatment for 18 hours was strongly potentiated by 75 µM HCQ and inhibited by higher concentrations. (*B*) HCQ increased LiCl-induced β-catenin signaling but at lower concentrations. (*C*) Macropinocytosis of TMR-dextran 70 kDa was stimulated by low doses of HCQ and inhibited at high levels. Note that in all cases HCQ was without effect in the absence of Wnt or LiCl treatment. Experiments represent biological triplicates. Error bars denote SEM (n ≥ 3) (**p < 0.01). (*D* and *D’*) HEK-293T cells have very low levels of macropinocytosis after 1 h of incubation with a macropinocytosis marker TMR-dextran 70 kDa. (*E* and *E’*) Adition of Wnt3a (100 ng/mL) increases macropinocytosis. (*F*-*I’*) Addition of HCQ potentiates Wnt-stimulated macropinocytosis, particularly at 75 µM HCQ. (*G* and *G’*) At higher concentration (100 µM), HCQ inhibits macropinocytosis. (*K* and *K’*) HCQ alone (75 µM) has no effect on macropinocytosis. Wnt3a-induced macropinocytosis and this is greatly potentiated by low-dose HCQ, inhibited by high dose HCQ, while HCQ alone has no effect on macropinocytosis The results of a similar experiment to that shown here were quantified by spectrophotometry in panel C. Scale bars 10 μm. See also Fig. S2.

Wnt3a triggers macropinocytosis of TMR-dextran 70 kDa (12) which has a hydrated diameter over 200 nm and is the gold standard for measuring macropinocytosis (22). Low-dose HCQ potentiated Wnt-induced macropinocytosis, while high doses inhibited it (Fig. 2*C*-*K*). In the absence of Wnt or LiCl, HCQ had no effect on macropinocytosis (Fig. 2*C* and *K’*). In animal cap explants, *xWnt8* mRNA strongly stabilized microinjected CD63-RFP, re-enforcing the view that the MVB compartment is expanded during Wnt signaling (*SI Appendix*, Fig. S2*M*-*N’*).

We then tested whether expansion of the MVB/lysosome compartment with HCQ could increase dorsalization in vivo. Microinjection of LiCl (4 nl at 300 mM) into a single ventral blastomere of the 4-cell *Xenopus* embryo resulted in slightly dorsalized embryos with enlarged head and cement gland structures (Fig. 3*A* and *B*). Co-injection of HCQ and LiCl resulted in a remarkable potentiation of dorsal development, generating embryos consisting mostly of head structures with reduced or absent trunks (Fig. 3*C* and 3*G*-*M*). This dorsalizing effect of HCQ was specific to LiCl and was not seen in NaCl controls (*SI Appendix*, Fig. S3*A*-*D*). While the effects of HCQ occurred over a very narrow concentration range in cultured cells, in the *Xenopus* embryo dorsalizing effects were found in the 1 to 10 mM range (in injections of 4 nl per embryo). At high HCQ levels (100 mM) no dorsalization was observed (*SI Appendix*, Fig. S3*E*-*H*).

**Fig. 3.**
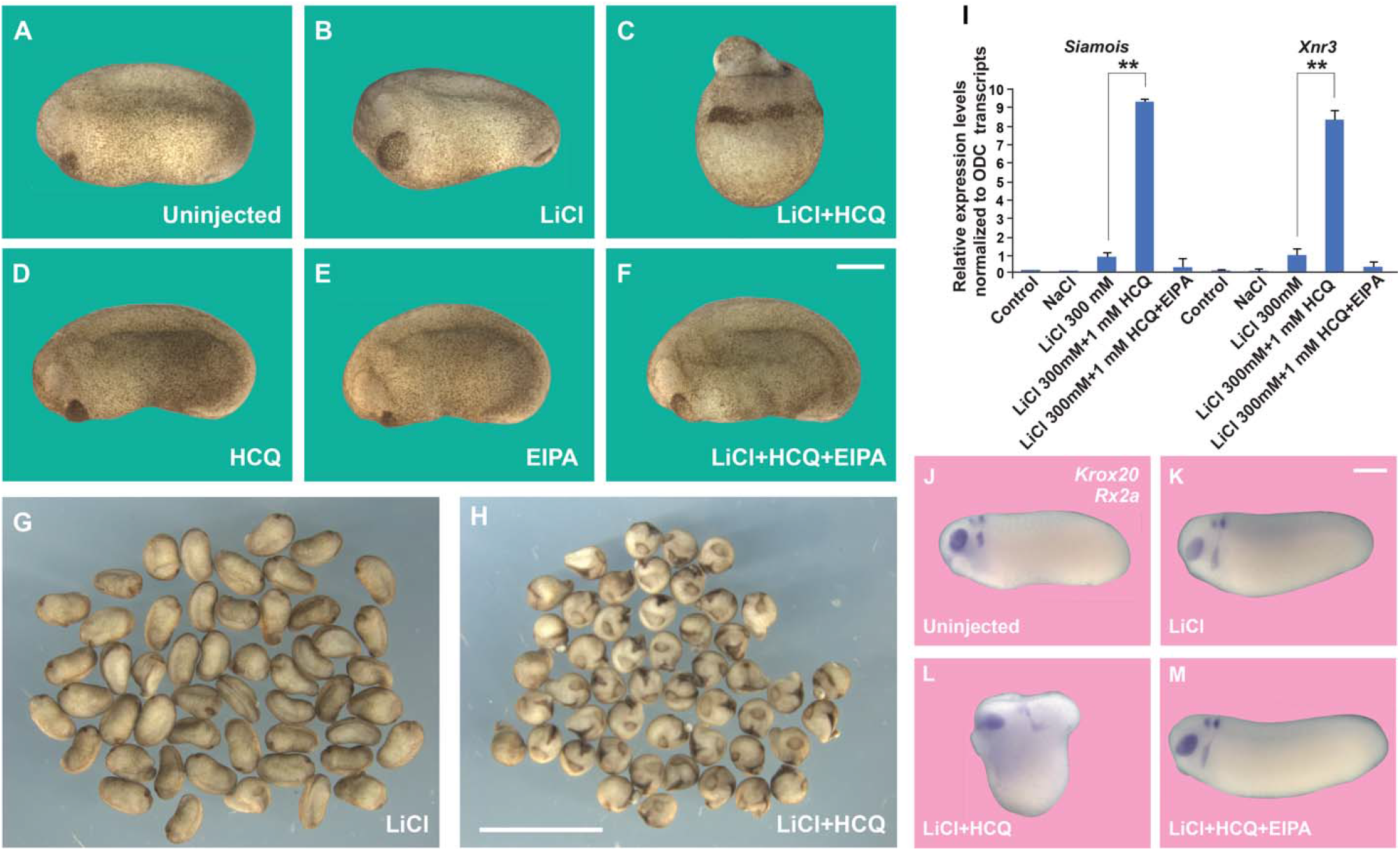
Dorsalization by LiCl Microinjection is Potentiated by HCQ in Xenopus Embryos Embryos were microinjected with 4 nl of 300 mM LiCl, 1 mM HCQ, 1 mM EIPA or their combinations one time ventrally at 4-cell stage. (*A*) Uninjected embryo at stage 24. (*B*) LiCl moderatly dorsalizes the embryo. (*C*) Low-dose HCQ strongly cooperates with LiCl. (*D* and *E*) HCQ and EIPA alone are without phenotype. (*F*) The macropincoytosis inhibitor EIPA blockes the dorsalization by LiCl plus HCQ. (*G* and *H*) panoramic comparison of LiCl alone and LiCl plus HCQ. (*I*) qRT-PCR at blastula stage 9.5 of the Wnt target genes Siamois and Xnr3 normalized for Ornithine decarboxylase (ODC). Note that HCQ strongly enhances the response to LiCl, that this is blocked by EIPA, and that NaCl serves as a control for LiCl. (*J*-*M*) In situ hybridization with Krox20 (hindbrain) and the Rx2a (eye) markers showing that the trunk is reduced and the head expanded in embryos dorsalized by LiCl plus HCQ, which was rescued by EIPA. Numbers of embryos analysed (5 independent experiments) were as follows A=182; B=137; C=174; D=142; E=115. Scale bars in F and K, 500 μm; Scale bar in H, 5 mm. Experiments in I represent biological triplicates; error bars denote SEM (n ≥ 3) (**p < 0.01). See also *SI Appendix*, Fig. S3.

EIPA is a Na^+^/H^+^ exchanger inhibitor derived from the diuretic Amiloride that blocks macropinocytosis through alkalinization of the cortical cytoplasm, preventing actin polymerization (22, 23). No phenotypic effect was observed with single injection of HCQ or EIPA (Fig. 3*D* and *E*). However, co-injection of 1 mM EIPA blocked the dorsalization phenotype caused by HCQ plus LiCl (Fig. 3, compare *C* to *F*). The potentiation of dorsal development by HCQ and LiCl takes place early in development, for it greatly increased expression of the cytoplasmic determinant target genes *Siamois* and *Xnr3* at the blastula stage, as determined by qRT-PCR (Fig. 3*I*). EIPA inhibited the increased *Siamois* levels (Fig. 3*I*) as well as V-ATPase subunit V0a3 stabilization (*SI Appendix*, Fig. S1). In addition, LiCl plus HCQ cooperated in stabilizing β-catenin protein on the dorsal side at blastula, and this was blocked by EIPA (*SI Appendix*, Fig. S3*I*-*N*). We conclude that low-dose HCQ increases activation of the Wnt signaling pathway at early stages of development and that this effect requires macropinocytosis.

### Lysosomes are Required for the Early Cytoplasmic Determinant Signal in *Xenopus*

We next tested whether lysosomes are required for activation of the dorsal early Wnt signal. The period of peak sensitivity of LiCl treatment has been carefully analyzed in *Xenopus* embryos and corresponds to the 32-cell stage (17). Our results indicated that dorsal lysosomes are already activated by LiCl at the 64-cell stage (Fig. 1); therefore, we titrated brief treatments of whole embryos by immersion in the lysosomal inhibitor Bafilomycin A1 at the 32-cell stage. Treatment with the V-ATPase inhibitor Baf at 5 µM for only 7 min before washing was found to be optimal. This brief treatment with Baf resulted in microcephalic embryos with no cement glands, in which the small head region was weakly pigmented while the ventro-posterior epidermis contained most of the maternal pigment (Fig. 4*A* and *B*). At early tailbud stage, the CNS marker *Sox2* was inhibited, reflecting the microcephaly (Fig. 4*C* and *D*). The ventralizing effect occurs early, for already at late blastula the effect of the brief treatment with the lysosomal acidification inhibitor Baf induced a sharp increase of the ventral development marker genes *Sizzled* (*Szl*) and *Vent1* by qRT-PCR (Fig. 4*E*, lanes 2 and 6). In LiCl-immersed embryos (300 mM, 7 min incubation at 32 cell) (17), the ventral markers *Szl* and *Vent1* were also strongly induced by subsequent treatment with Baf (Fig. 4*E*, lanes 4 and 8). Immersion in LiCl resulted in radial dorsal tadpole stage embryos with increased pan-neural Sox2 expression surrounding the blastopore, while subsequent incubation with Baf at the 32-cell inhibited CNS induction (*SI Appendix*, Fig. S4*A*-*C’*). The dominant effect of the lysosomal acidification inhibitor Baf is in agreement with the increased membrane trafficking and lysosomal acidification elicited by Wnt and LiCl in mammalian cultured cells (10).

**Fig. 4.**
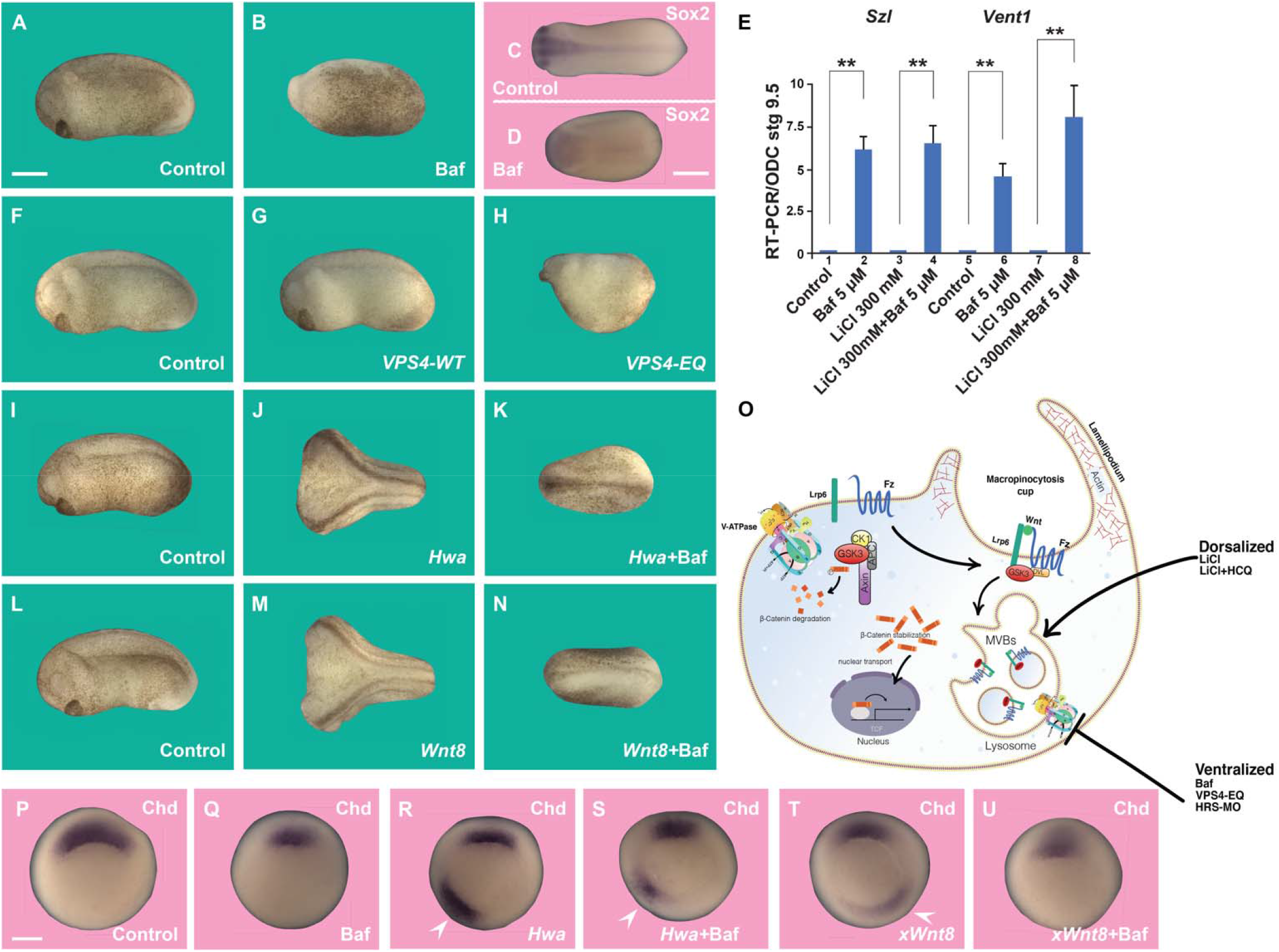
Dorsal Cytoplasmic Determinant Activity Requires Lysosomal Acidification and the ESCRT Machinery; Hwa Signaling is Inhibited by Bafilomycin. (*A* and *B*) Immersion of *Xenopus* embryos at 32-cell stage in 5 µM Baf for 7 min results in a partially ventralized phenotype with microcephaly, loss of cement gland and pigmentation in ventral and posterior ectoderm. (*C* and *D*) Baf reduces the pan-neural marker *Sox2*. (*E*) qRT-PCR of blastula embryos showing that Baf increases expression of the ventral markers Szl and Vent1 in wild-type and LiCl-treated embryos. (*F*-*H*) Inhibiting ESCRT machinery with Vps4-EQ, but not Vps4-WT, mRNA (two dorsal injections of 400 pg at 4-cell stage) reduces axis formation. (*I*-*K*) Microinjection of *xHwa* mRNA (10 pg one time ventrally) induces complete secondary axes that are blocked by Baf incubation at 32 cell stage. (*L* and *N*) *xWnt8* mRNA second axes are also inhibited by the V-ATPase inhibitor Baf. (*O*) Model of membrane trafficking during Wnt signaling highlighting macropinocytosis, V-ATPase, MVBs, and lysosomes. (*P*-*U*) In situ hybridizations of Chd at the gastrula stage showing partial inhibition of the organizer by Baf in control, Hwa and xWnt8 axes; arrowheads indicate ectopic *chd* expression. Numbers of embryos analysed were as follows A=94; B=77; F=89; G=62; H=46; I=235; J=67; K=71; L=182; M=57; N=70 (2 independent experiments). Scale bars 500 μm. qRT-PCR experiments represent biological triplicates, error bars denote SEM (n ≥ 3) (**p < 0.01). See also *SI Appendix*, Fig. S4.

Wnt signaling requires endocytosis and the sequestration of GSK3 in MVB/endolysosomes via the ESCRT machinery (9, 10). Vacuolar protein sorting 4 (Vps4) is an ATPase essential for the final steps of MVB intraluminal vesicle formation and has a very effective dominant-negative point mutation (Vps4-EQ) that blocks its function (24). Microinjection of Vps4-EQ mRNA into each dorsal blastomere at the 4-cell stage reduced axial development and resulted in microcephalic embryos and, importantly, this inhibition was specific since Vps4-WT had no effect on the embryo (Fig. 4*F*-*H*). Microinjection of the validated antisense reagent Hrs-MO (9) also inhibited anterior and dorsal development (*SI Appendix*, Fig. S4*D*). These results indicate that dorsal lysosomes and components of the MVB membrane trafficking machinery are required for the full activity of the endogenous dorsal signal in *Xenopus* embryos.

Hwa is a maternal mRNA encoding a novel transmembrane protein translated specifically in dorsal blastomeres in zebrafish and *Xenopus* embryos. It is likely that Hwa is the previously mysterious dorsal cytoplasmic determinant activated after fertilization (5). When microinjected into a ventral blastomere at 4-8 cell stage, *Hwa* mRNA induced complete secondary axes with high efficiency (Fig. 4*E* and *J*). This result provides confirmation by an independent laboratory that *xHwa* mRNA has potent inductive effects (5). Since dorsal development required lysosomal acidification, we also tested whether Baf treatment inhibited Hwa function. We found that 7 min incubation of 32-cell embryos with the V-ATPase inhibitor Baf strongly inhibited secondary and primary axis formation in Hwa-injected embryos (Fig. 4*J* and *K*). Co-injection of Hwa mRNA with EIPA or Hrs-MO blocked Hwa inductive activity, further strengthening the view that Hwa requires the membrane trafficking machinery (*SI Appendix*, Fig. S4*E*-*H*). We also examined the effect of Baf in *xWnt8* mRNA injected embryos, in which the same signaling pathway is activated. It was found that Baf blocked the Wnt8 twinning phenotype (Fig. 4*L*-*N*). Baf treatment partially inhibited expression of the Spemann organizer gene Chordin (Chd) in endogenous, Hwa and xWnt8 axes (Fig. *P*-*U*). Taken together, the results indicate that lysosome activity and MVB formation is required for Hwa signaling at the 32-cell stage.

## DISCUSSION

The main conclusion of the present study is that lysosomes play an essential role in the establishment of the initial polarity of the body axis in *Xenopus* embryos. As indicated in Fig. 4*O*, recent work has implicated membrane trafficking as a key step in the activation of Wnt signaling (10). After fertilization, a microtubule-driven cortical rotation of the egg generates the first asymmetry in the egg, resulting in a dorsal Wnt-like signal (Fig. 1*A’*) (3, 4). Wnt signaling causes a remarkable increase in macropinocytosis, as well as lysosomal activation and acidification, even in the absence of new protein synthesis (12, 13). As the role of lysosomes in development was largely unexplored, we examined the localization of active lysosomes at the 64-cell stage, just after the period of peak sensitivity to GSK3 inhibition by LiCl (17). Lysosomes containing activated cathepsin D were enriched in dorsal marginal cells, particularly in embryos microinjected with Wnt or LiCl (Fig. 1). Involvement of the MVB/lysosome compartment was suggested by the marked increase in dorsalization by LiCl at low doses, but not high doses, of HCQ that expanded the MVB compartment marked by CD63. The transit through intraluminal vesicles of MVBs is a required step for the trafficking of plasma membrane protein into lysosomes (10). A pulse of the lysosome acidification inhibitor Bafilomycin A1 at 32-cell stage increased ventral markers and decreased endogenous Spemann organizer formation, as did interfering with the ESCRT machinery that forms MVBs.

In mammalian cells, low-dose HCQ increased Wnt signaling but only within narrow ranges of concentration. It is conceivable that some of the clinical prophylactic antimalarial and anti-autoimmune effects of HCQ might be mediated by increased macropinocytosis and lysosomal activity in small populations of Wnt-activated immune cells, rather than lysosomal inhibition as is currently thought (25 26).

The recent discovery of the maternal Hwa transmembrane protein has suggested that after fertilization in dorsal cells Hwa, rather than a canonical Wnt-Lrp6 interaction, is responsible for the dorsal maternal signal (5, 27). Hwa has is a plasma membrane protein with a short extracellular domain and a large intracellular domain that functions by promoting the degradation of Axin1 in zebrafish and *Xenopus* embryos (5). In a mammalian cell system, zebrafish Hwa binds to Axin1 and promotes its degradation with the involvement of tankyrases (5). The tumor suppressor Axin1, which is a GSK3-binding protein, is a component of the β-catenin destruction complex and its degradation stabilizes β-catenin, leading to the transcriptional activation of the Wnt signaling pathway (6, 27). Importantly, a recent report from the Tao laboratory showed that the dorsalizing activity of Hwa is opposed by ubiquitination via the transmembrane E3 ubiquitin ligase zinc and ring finger 3 (ZNRF3) and trafficking into lysosomes to limit or dampen Spemann organizer formation in the ventral side of the embryo (28). At first glance, these results appear to contradict our present findings that lysosome activity and MVB formation are required for signaling by microinjected *Hwa* mRNA and endogenous axis formation. However, the results presented here are concerned with brief 32-cell stage Baf treatments which should target the preferentially the lysosomes that were found to be activated on the dorsal side of the marginal zone at the 64-cell stage. Interestingly, in ref. 28 experiments using embryos derived from oocytes microinjected with HA-tagged Hwa mRNA displayed Hwa-HA protein stabilization by Baf treatment at the 32-cell stage (3 hours of development). This suggests that the lysosomal regulation of Hwa can take place at the stages when Hwa is promoting dorsal development. Work from our laboratory indicates that the lysosomal membrane trafficking pathway is a required step in the canonical Wnt signaling pathway (12, 13). We propose that on the dorsal side of the early embryo non-ubiquitinated Hwa may bind Axin1, triggering its trafficking into lysosomes together with GSK3, resulting in the sequestration of GSK3 to generate a Wnt-like signal. In future, it will be interesting to investigate whether endogenous Hwa protein colocalizes with GSK3 in dorsal MVBs/lysosomes in early *Xenopus* embryos.

How does Wnt signaling regulate lysosome activation? Previous work has shown the activated Wnt Lrp6 receptor binds to the peripheral V-ATPase protein ATP6AP2 (also called Prorenin receptor) (8). ATP6AP2 binds to transmembrane protein 9 (TMEM9), which in turn binds to and activates the V-ATPase that acidifies lysosomes (29). TMEM9 is an MVB protein that induces secondary axes in *Xenopus* embryos and plays an oncogenic role in colorectal (CRC) and hepatocellular (HCC) carcinomas (30). Treatment with Baf reduces tumor growth in TMEM9-expressing CRC and HCC xenografts, indicating an important role for lysosomes in tumor progression (29). It is possible that in the *Xenopus* embryo V-ATPase and lysosomal trafficking is activated on the dorsal side by a similar TMEM9-V-ATPase activation. In sum, membrane trafficking, macropinocytosis, V-ATPase, and lysosomes are essential for the earliest developmental decisions in the *Xenopus* vertebrate development model system.

## Materials and Methods

### Reagents and Antibodies

SiR-Lysosome was purchased from Cytoskeleton (CYSC012, used at 1 μM). Hydroxychloroquine (H0915), Chloroquine diphosphate (C 6628), 5-(N-Ethyl-N-isopropyl) amiloride (EIPA) (A3085), LiCl and Sodium chloride (NaCl) (S9888), IPA-3 (I2285, used at 2.5 μM) were obtained from Sigma. Bafilomycin A1 (Baf-A1) (S1413) was purchased from Selleckchem. Tetramethylrhodamine Dextran (TMR-Dx) 70,000 kDa was purchased from ThermoFisher (D1818). Total β-catenin antibody (1:1000) was purchased from Invitrogen (712700), GAPDH (1:1000) was obtained from cell signaling, anti-ATP6V0a3 antibody (*23*) was obtained from Novus (nbp1-89333 1:1000). Antibodies against Pak1 (ab131522), Ras (ab52939), and secondary antibodies for immunostaining (ab150083, ab150117) (1:500) were obtained from Abcam. Wnt3a protein was from Peprotech (315-20) and used at 100 ng/ml. HRS-MO TGCCGCTTCCTCTTCCCATTGCGAA (*9*) was from Gene Tools and microinjected as 4 nl of 0.3 mM MO.

### Tissue Culture

HEK-293BR (BAR/Renilla) were cultured in DMEM containing 10% fetal bovine serum (FBS), SW480 cells and SW480APC cells (31) were cultured at 37 ° C in 5% CO2 atmosphere in DMEM/F12 (Dulbecco’s Modified Eagle Medium: Nutrient Mixture F-12), supplemented with 5% fetal bovine serum, 1% glutamine, and penicillin/streptomycin. The cells were seeded at a cell density of 20%-30% and experiments were performed when cells reached a confluence of 70-80%. Cells were cultured 6-8 hours in medium with 2% fetal bovine serum before treatments.

### *Xenopus* Embryo Microinjection and In Situ Hybridization

*Xenopus laevis* embryos were fertilized *in vitro* using excised testes and cultured and staged as described (32). Ventral co-injection 4 nl LiCl (300 mM) ± HCQ (1-2 mM) was performed 2-4 cell stage and cultured until early tadpole stage. EIPA was injected at 1 mM. pCS2+ clones encoding *xWnt8*, x*Hwa*, CD63-RFP, mGFP, Vps4-WT and Vps4-EQ, were linearized with NotI and transcribed with SP6 RNA polymerase using the Ambion mMessage mMachine kit. Embryos were injected in 1 x MMR saline and cultured in 0.1 x MMR. *In vitro* synthesized mRNAs were microinjected in a 4 nl volume using an IM 300 Microinjector (Narishige International USA, Inc). To study the role of ESCRT in development, *Xenopus* antisense HRS MO (Taelman et al., 2010) was injected two times dorsally at the 4-cell stage. In Situ hybridizations were performed as described previously at http://www.hhmi.ucla.edu/derobertis.

### Bisected embryo lysosome staining

Embryos at 2-4 cell stage were injected once into a ventral blastomere with LiCl (300 mM) or *xWnt8* (2 pg). Embryos were incubated in 0.1 x MMR solution until 64-cell stage. Embryos were collected and fix with 4% PFA (paraformaldehyde, Thermo Scientific, 28908.) diluted in 1 x PBS (Fisher Scientific, BP399-4). After 24 h of fixation, embryos were cut along the D-V axis with a surgical blade and re-fixed with 4% PFA for 15 min. Embryos were then washed with PBS three times, blocked with 5% BSA in PBS to reduce background for 24 h, and incubated with 1 μM Sir-Lysosome marker for active lysosomal marker cathepsin D for 2 hrs, washed 3 times with PBS. Samples were imaged using an Axio Zoom.V16 Stereo Zoom Zeiss microscope with Apotome function and stacked images reconstructed using the Zen 2.3 pro Zeiss software. For embryo stainings with V-ATPase subunit V0a3 or β-catenin antibodies, after the BSA blocking treatment samples were incubated with the primary antibody (1:300) overnight, washed, then incubated with the secondary antibody also overnight (1:500). Embryos were plated in agar Petri dishes with PBS for microscopic examination as soon as possible to avoid the loss of fluorescence.

### Animal Cap Cell Culture

Animal caps were dissected at early blastula stage 8.5 to 9 in 1 x MMR solution, washed 3 times, and cells cultured in L-15 medium containing 10% Fetal Calf Serum diluted to 50% with H_2_0 (33) on fibronectin-coated 35 mm Petri dishes with glass bottoms (#0 cover glass, Cell E&G: GBD00003-200) for 12-18 h. When mounting with coverslips, a drop of Anti-fade Fluorescence Mounting Medium-Aqueous, Fluoroshield (ab104135) was added and microscopic examination of RFP and GFP performed. Image acquisition was performed using a Carl Zeiss Axio Observer Z1 Inverted Microscope with Apotome.

### Immunostainings of mammalian cells

HEK293T and SW480 ± APC cells were plated on glass coverslips. Coverslips were acid-washed and treated with 10 μg/ml Fibronectin (Sigma F4759) for 30 min at 37°C to enhance cell spreading and adhesion. Cells were fixed with 4% paraformaldehyde (Sigma #P6148) for 15 min, permeabilized with 0.2% Triton X-100 in PBS for 10 min, and blocked with 5% BSA in PBS for 1 hour. Primary antibodies (1:100) were added overnight at 4°C. The samples were washed three times with PBS, and secondary antibodies were applied for one hour at room temperature (1:1000). After three additional washes with PBS, coverslips were mounted with Fluoroshield Mounting Medium with DAPI (ab104139). Immunofluorescence was analyzed and photographed using a Zeiss Imager Z.1 microscope with Apotome.

### TMR Dextran Assay

HEK293T cells were plated on glass coverslips for 12-18 h. Cells were then incubated at 37°C in 5% CO2 atmosphere with DMEM supplemented with 2% fetal bovine serum, and 1 mg/ml TMR for one hour. After incubation, cells were washed, fixed and blocked for 1 h with 5% BSA in PBS to reduce background from nonspecific binding. The coverslips were mounted with Fluoroshield Mounting Medium with DAPI (ab104139). Immunofluorescence was analyzed and photographed using a Zeiss Imager Z.1 microscope with Apotome and using the Zen 2.3 imaging software.

### Western Blots and Luciferase Assays

Cell lysates were prepared using RIPA buffer (0.1% NP40, 20 mM Tris/HCl pH 7.5), 10% Glycerol, together with protease (Roche #04693132001) and phosphatase inhibitors (Calbiochem #524629) and processed as described (12). Stable cell line HEK-293-BR expressing firefly luciferase genes (12) were incubated for 12-18 h with Wnt3a (100 ng/ml.) or LiCl (40 mM) plus or minus HCQ (15-90 μM) or with HCQ alone. Luciferase activity was measured with the Dual-Luciferase Reporter Assay System (Promega) according to manufacturer’s instructions, using a Glomax Luminometer (Promega).

### Measurement of TMR uptake by Spectrophotometry

Cells were seeded to maintain a cell density between 20%-30% and experiments were performed when cells reached a confluence of 70-80%. Cells were starved with 2% BSA for 6-12 h, then stimulated with Wnt3a (100 ng/ml) plus or minus HCQ (15-100 μM) for 12-18 h in 96-well plates. After incubation, TMR-dextran 70 kDa was added to the medium for 1 h (1 mg/ml). Cells were washed 3 times with PBS, trypsinized and transferred into a 96 well microplate for reading. The TMR-dextran 70 kDa (Excitation/Emission 555/580 nm) absorbance was quantified with a SpectraMax microplate reader using the Softmax pro 5.4.4 software. The conditions used for Excitation/Emission were 544/590 nm. Three independent measurements were performed for each sample in this quantitative assay.

### Bafilomycin Treatments

Frog embryos at 32-cell stage were incubated with Bafilomycin A1 (Baf, 5 μM) for 7 min in 0.1 MMR solution. After incubation, embryos were washed 2 times with 0.1 MMR solution and cultured overnight until early tailbud tadpole stage. Embryos injected once ventrally at 4-cell stage with *xWnt8* (2 pg) or *Hwa* (10 pg) mRNA were incubated in 0.1 x MMR solution until 32-cell stage. Embryos were then transferred in 0.1 x MMR solution containing Baf (5 μM) and incubated for 7 min. After that, the embryos were washed 2 times with 0.1 x MMR solution and cultured until tadpole stage. For whole embryo LiCl treatments 32-cell stage embryos were incubated with LiCl (300 mM) for 7 min. After that, embryos were quickly washed two times with 0.1 MMR solution and then incubated for 7 min in 0.1 MMR solution containing 5 μM Baf. After the treatment, the embryos were washed 2 times with 0.1 MMR solution and cultured until tadpole stage.

### qRT-PCR

Quantitative RT-PCR experiments using *Xenopus* embryos were performed as previously described (32). Primer sequences for qRT-PCR are as follows:

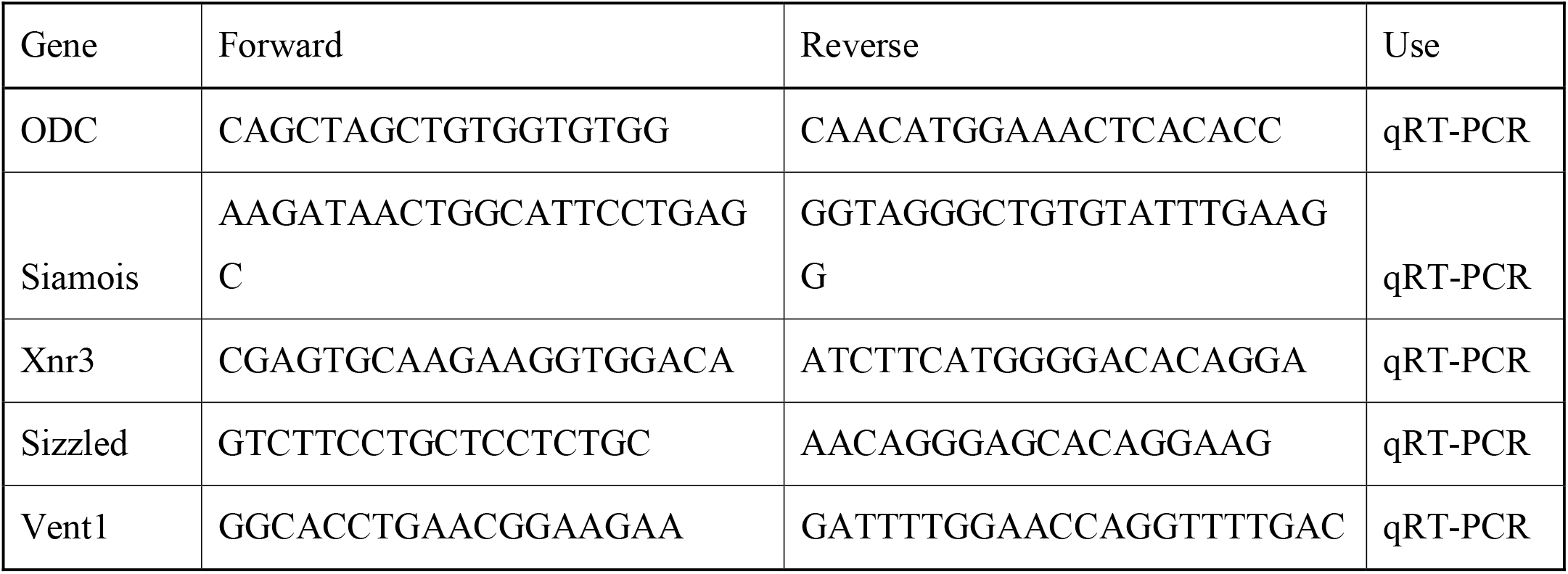

### Statistical Analyses

Data were expressed as means and standard errors of the mean (SEM). Statistical analysis of the data was performed using the Student t test; a P-value of <0.01** was considered statistically significant for differences between means.

### Data Availabilty

All study data are included in the article and/or *SI Appendix*.

## ACKNOWLEDGEMENTS

We are grateful to J. Monka and D. Geissert for technical assistance, Q. Tao for xHwa construct, R. Moon for BAR reporters and xWnt8, M. Faux for SW480APC cells, Y. Moriyama, Y. Ding, and J. Monka for comments on the manuscript, and M. F. Domowicz for help with the illustrations. This work was supported by the UC Cancer Research Coordinating Committee (grant C21CR2039); National Institutes of Health grant P20CA016042 to the University of California, Los Angeles Jonsson Comprehensive Cancer Center; and the Norman Sprague Endowment for Molecular Oncology.

## Supplemental Information Appendix

**SI Appendix,Fig. S1.**
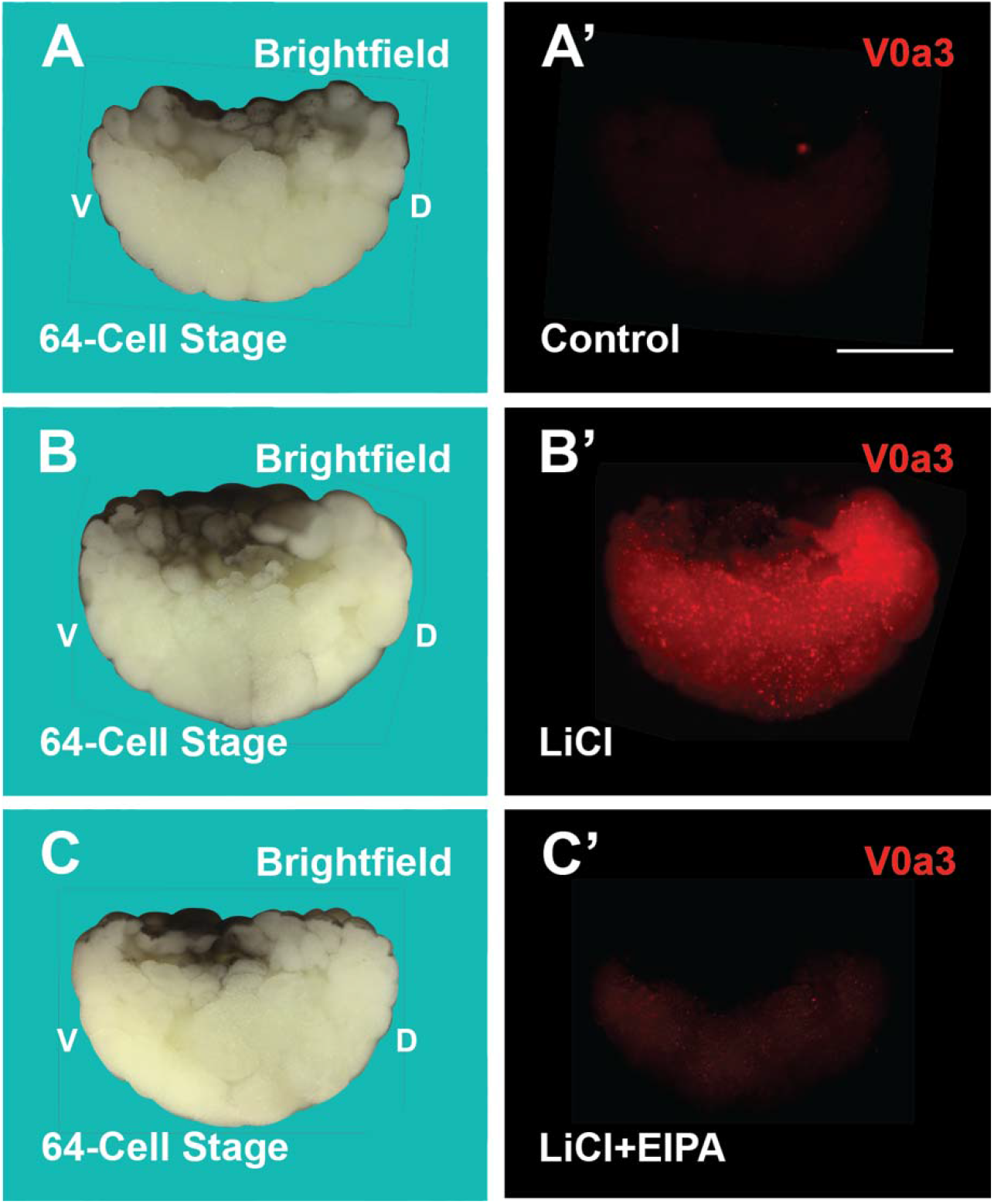
Microinjection of LiCl at 2-4 cell-stage Stabilized the V-ATPase Subunit V0a3, Particularly in Dorsal Cytoplasm, at the 64-cell Stage, and this was Blocked by the Macropinocytosis Inhibitor EIPA. Fixed embryos were bisected and the dorsal (D) and ventral (V) sides determined by their pigmentation and thickening of the dorsal marginal zone. The anti-V0a3 antibody has been validated (18). (*A* and *A’*) Uninjected bisected embryos. (*B* and *B’*) LiCl microinjection strongly stabilizes the lysosomal V-ATPase subunit V0a3, especially on the dorsal side. (*C* and *C’*) Co-injection of 1 mM EIPA, a micropinocytosis inhibitor prevents V-ATPase stabilization. Embryos analysed were as follows A, n=27; B, n=30; C, n=32 half-embryos with D-V polarity. Scale bars 500 μm.

**SI Appendix,Fig. S2.**
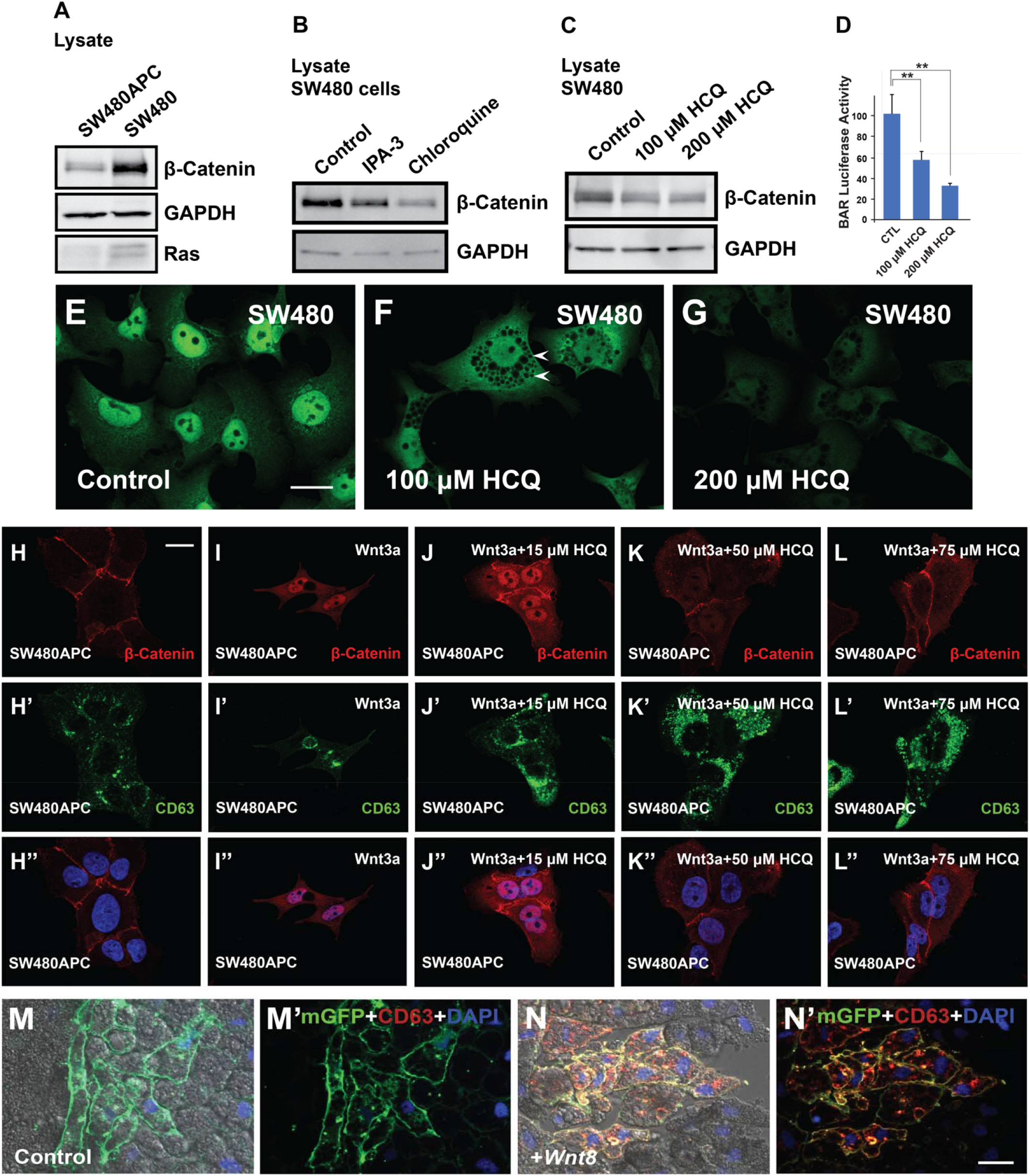
CQ and HCQ at High Concentrations Inhibit the Constitutive Wnt Signaling Present in Colorectal Carcinoma SW480 Cells in which the Tumor Suppressor APC is Mutated; when APC is Reconstituted in SW480-APC Cells, Low-dose HCQ Expands the CD63-positive MVB Compartment and Nuclear β-catenin Levels, whereas at High Doses it Inhibits β-catenin Accumulation while Retaining Increased CD63 Endosomes; in *Xenopus* Ectodermal Explants Wnt Signaling Causes the Stabilization of CD63-RFP Protein. (*A*) SW480 cells, but not SW480APC cells, have stabilized β-catenin and total Ras. GAPDH serves as a loading control. (*B*) CQ (250 µM) or the Pak-1 inhibitor IPA-3 (2.5 µM) decreases β-catenin levels in SW480 cells. Pak-1 is a protein kinase required for macropinocytosis in Wnt signaling (10). (*C*) HCQ at high concentrations inhibits constitutive β-catenin levels in SW480 cells. (*D*) HCQ inhibits β-catenin transcriptional activity. (*E*) SW480 cells have very strong constitutive nuclear β-catenin staining. (*F*) HCQ treatment at 100 µM for 12 hours reduces β-catenin levels and result in greatly enlarged endolysosomal compartment vesicles (arrowheads). (*G*) HCQ at 200 µM for 12 hours significantly reduced β-catenin levels. These results suggest that lysosome inhibition could provide a therapeutic strategy for cancers caused by mutations in tumor suppressors in the Wnt pathway. (*H*-*H’’*) Colorectal carcinoma SW480 cells reconstituted with APC (31) have low levels of nuclear β-catenin and CD63 MVBs. (*I*-*I’’*) SW480APC cells respond to Wnt3a by accumulating nuclear β-catenin. (*J*-*J’’*) 15 μM HCQ causes the accumulation of CD63 MVBs and β-catenin. (*K*-*L’’*) Higher concentrations of HCQ inhibit β-catenin nuclear accumulation but retain CD63 endolysosomes. Note that different cell lines differ in the optimal concentration of HCQ to potentiate Wnt or LiCl. (*M*-*M’*) Animal cap cells co-injected with *membrane GFP* (*mGFP*) and *CD63-RFP* mRNA. A single animal pole cell was injected at 8-cell stage, ectodermal explants excised at late blastula, attached to fibronectin glass chambers and cultured overnight in L-15 medium. (*N*-*N’*) Co-injection of *xWnt8* mRNA results in the stabilization of CD63-RFP, which accumulates in cytoplasmic membrane vesicles. Scale bars 10 μm.

**SI Appendix,Fig. S3.**
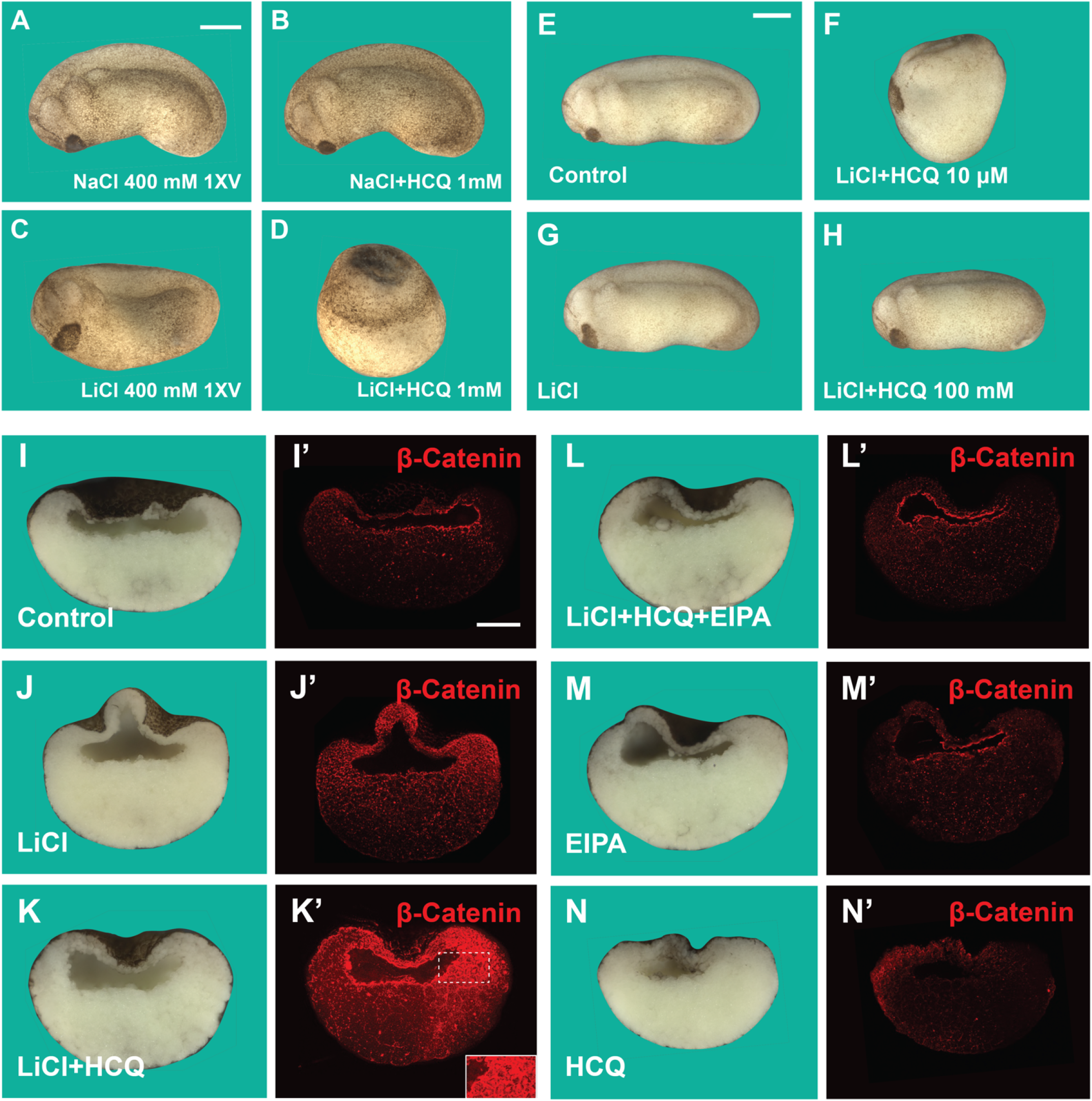
LiCl, but not NaCl, Cooperated with HCQ in the 1-10 mM Range while 100 mM HCQ did not Increase Dorsalization; β-catenin Protein is Stabilized by Microinjection of LiCl, Potentiated by Co-injection of HCQ, and Blocked by EIPA at Blastula Stage. (*A*) Embryo injected with 400 mM NaCl has normal development at stage 24. (*B*) NaCl does not cooperate with HCQ, providing a specificity control. (*C*) Embryo weakly dorsalized by LiCl alone (4 nl at 300 mM ventrally). (*D*) LiCl plus HCQ (1 mM) showing a completely radial dorsalized phenotype with circular cement gland. (*E* and *F*) 10 mM LiCl cooperates with HCQ; note that the *Xenopus* embryo has a wider concentration range for the HCQ effect in comparison to cultured mammalian cells. (*G* and *H*) At a high concentration of (100 mM) HCQ does not cooperate with LiCl, as is the case in cultured cells. (*I* and *I’*) Endogenous β-catenin was almost undetectable in control embryos at blastula stage. (*J* and *J’*) β-catenin protein was stabilized by injection of the GSK3 inhibitor LiCl. (*K* and *K’*) Dorsal β-catenin was greatly enhanced by co-injection of HCQ and LiCl; inset shows both nuclear and cell membrane localization. (*L* and *L’*) The macropinocytosis inhibitor EIPA (1 mM) blocked the effects of LiCl plus HCQ. (*M*-*N’*) EIPA or HCQ alone do not affect β-catenin protein staining. These results confirm that LiCl plus HCQ are strongly dorsalizing and that Na^+^/H^+^ exchanger inhibition blocks this effect. Numbers of embryos analysed were as follows A=52; B=58; C=60; D=78; E=51; F=28; G=29; H=45. In I-N’ about n=30 half-embryos were analysed per condition. Scale bar 500 μm.

**SI Appendix,Fig. S4.**
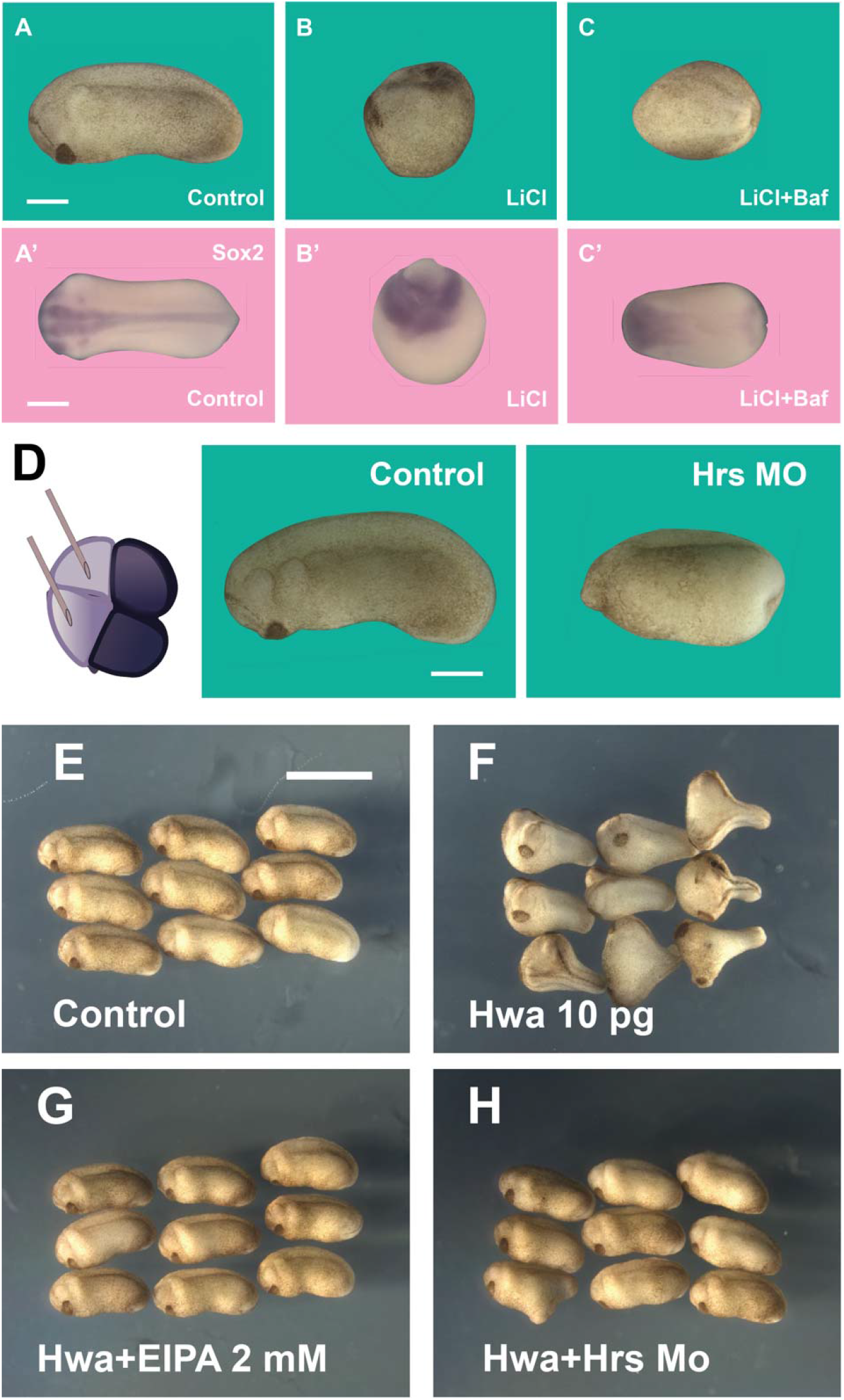
LiCl Treatment of Embryos Radially Dorsalized Embryos and this was Blocked by Subsequent Inhibition of V-ATPase; Twinning by Microinjected Cytoplasmic Determinant Hwa Requires Membrane Trafficking. (*A*) Control embryo cultured in 0.1 MMR saline. (*B*) Embryo partially dorsalized by LiCl (300 mM for 7 min), note the blastopore at the top and lack of trunk-tail structures. (*C*) Embryo treated with LiCl and subsequently immersed in Baf (5 µM for 7 min) in 0.1 MMR saline; note that LiCl effect is blocked. (*A’*-*C’*) In situ hybridizations of siblings with the pan-neural marker Sox2. Note the radial CNS surrounding the blastopore and that Baf blocks the increased neural signal, consistent with the microcephaly phenotype observed after treatment of WT embryos. (*D*) Microinjection of antisense Hrs MO, known to specifically block MVB formation (9), interferes with dorso-anterior development. Embryos were injected with MOs 2 times dorsally at the 4-cell stage. Note the lack of head and cement gland, indicating a requirement for the ESCRT machinery. (*E*) Uninjected controls for Hwa experiment. (*F*) Hwa mRNA induces 100% of complete secondary axes. In our hands, Hwa is even more effective than Wnt in inducing complete twinning. (*G*) Co-injection of the macropinocytosis inhibitor EIPA abolishes Hwa activity. (*H*) Interfering with MVB formation by co-injection of Hrs MO (0.3 mM, 4 nl) prevents Hwa function. Together with the inhibition of Hwa by Baf (Fig. 9*I*-*K* in main text), this results strongly suggest that Hwa transmembrane protein activates endocytosis through the Wnt signaling pathway. Numbers of embryos analysed were A=76; B=110; C=109 per condition, 4 independent experiments; in D Controls were n=115, HRS MO n=107, 7 independent experiments; E=148; F=142; G=120 (3 independent experiments); H=45 per condition. Scale bars for A and D, 500 μm, for E, 2 mm.

